# Food intake dependent and independent effects of heat stress on lactation and mammary gland development

**DOI:** 10.1101/2020.04.03.024679

**Authors:** Yao Xiao, Jason M. Kronenfeld, Benjamin J. Renquist

## Abstract

With a growing population, a reliable food supply is increasingly important. Heat stress reduces livestock meat and milk production. Genetic selection of high producing animals increases endogenous heat production, while climate change increases exogenous heat exposure. Both sources of heat exacerbate the risk of heat-induced depression of production. Rodents are valuable models to understand mechanisms conserved across species. Heat exposure suppresses feed intake across homeothermic species including rodents and production animal species. We assessed the response to early-mid lactation or late gestation heat exposure on milk production and mammary gland development/function, respectively. Using pair-fed controls we experimentally isolated the food intake dependent and independent effects of heat stress on mammary function and mass. Heat exposure (35°C, relative humidity 50%) decreased daily food intake. When heat exposure occurred during lactation, hypophagia accounted for approximately 50% of the heat stress induced hypogalactia. Heat exposure during middle to late gestation suppressed food intake, which was fully responsible for the lowered mammary gland weight of dams at parturition. However, the impaired mammary gland function in heat exposed dams measured by metabolic rate and lactogenesis could not be explained by depressed food consumption. In conclusion, mice recapitulate the depressed milk production and mammary gland development observed in dairy species while providing insight regarding the role of food intake. This opens the potential to apply genetic, experimental and pharmacological models unique to mice to identify the mechanism by which heat is limiting animal production.

**Summary Statements:** This study demonstrates that heat stress decreases lactation and mammary development through food intake dependent and independent mechanisms.

## INTRODUCTION

Rising global temperatures may result in global food insecurity. Heat exposure in livestock species decreases feed intake, depressing meat and milk production. Heat abatement strategies, which are largely restricted to intensive production systems limit the losses associated with heat stress. Despite implementation of heat abatement, in 2003 production losses cost $897 million, $369 million, $299 million, and $128 million for the U.S. dairy, beef, swine, and poultry industries, respectively (St-Pierre et al., 2003). By 2019, the dairy industry’s annual heat induced economic loss had risen to $1.2 billion (Key and Sneeringer, 2014). This cost is predicted to keep rising throughout the 21^st^ century with climate change (Gunn et al., 2019). Although the economic costs are robust, the water intensive nature of most heat abatement strategies exacerbates the environmental impact of animal production. With water shortages common to many areas and environmental concerns of agriculture runoff, the development of approaches to limit production losses while restricting water use is essential.

The economic losses of billions of USD are due to depressed growth and milk production. Pair-feeding studies establish that across species the heat induced depression in growth is nearly entirely attributable to hypophagia (O’Brien et al., 2010; Zeferino et al., 2016; Zhao et al., 2018). Thus, by understanding the mechanism behind heat stress hypophagia we may be able to restore growth. The decrease in milk production that results from heat exposure is multi-faceted. It can be broken into short-term hypogalactia or depressed mammary development and hypophagia independent or dependent. In short-term hypogalactia 50% of the decrease in milk yield is a response to hypophagia (Rhoads et al., 2009; Wheelock et al., 2010). Heat stress during the critical stages of mammary gland development (prior to and immediately following parturition) dramatically affects mammary gland development and milk production throughout lactation (Collier et al., 1982; Tao et al., 2011). The role of decreased energy intake on this muted mammary gland development has not been isolated from possible hypophagia independent effects.

Rodents display a heat induced reduction in feed intake that mimics that in production animal species (Hepler et al., 2016; Huynh et al., 2005; Lu, 1989; Morera et al., 2012; Spiers et al., 2004). Moreover, rodents recapitulate the increased sensitivity to elevated external temperature during lactation and gestation observed in high producing dairy cows (Gantner et al., 2017; Simons et al., 2011; Tao et al., 2011). With the conservation of phenotype, the genetic, pharmacological, and surgical models available in rodents may open the door for research aimed at understanding the mechanism by which heat depresses food intake, milk production, and late gestation mammary development. Accordingly, we report the development of mouse models to assess the food intake dependent and independent effects of heat on milk production and mammary development.

## MATERIALS AND METHODS

### Animals

Male and female C57BL/6J mice were purchased from The Jackson Laboratory (Bar Harbor, ME) and singly housed so that we could assess individual food intake. Mice were maintained in a 14h light, 10h dark light cycle. All experimental protocols were approved by the Institutional Animal Use and Care Committee at the University of Arizona.

### Food intake experiment

Control mice were housed at the control (CTL) environment (22°C, 50% relative humidity) and given ad libitum access to NIH-31 chow (Harlan Laboratories, Indianapolis, IN) and water. Food and water weights were measured at 0600 h and 1800 h daily to measure day and night consumption. Heat stress (HS) mice were placed in a heat chamber (Coy Lab Products, Grass Lake, MI) set at 35°C and 50% humidity. These environmental settings have been shown to suppress food intake in mice previously (Hepler et al., 2016; Morera et al., 2012).

### Lactation experiment

Eight-month-old multiparous females were used for these studies to ensure that mammary gland development was not limiting milk production. Within one day post-parturition, litters were culled to 6 pups. Dams with a litter of only 5 pups remained on the study, while those with smaller litters were culled. Pair-fed (PF) mice were housed in the control environment, but fed to match food consumed by HS mice. To match diurnal food consumption patterns, ∼25% of daily feed allotment was provided to PF mice at 1000 h and ∼75% at 1800 h. Food intake, water intake, body mass and litter mass were recorded daily at 1000 h from 4 to 11 days postpartum (dpp). Treatments (CTL, HS or PF) began at 5 dpp. The weigh-suckle-weigh method was employed to assess milk production (Hernandez et al., 2012). Briefly, pups were separated from dams for 4 h (1000 h to 1400 h). At the end of this 4 h separation, the litter was weighed and transferred back into the original cages with the dam. After 1h, the litter was again weighed. The litter mass gain during the 1 h of suckling acted as a proxy of dam milk production.

### Mammary gland experiment

We used 4 months old virgin females to assess the effect of heat stress on mammary gland development. To time breeding, 2∼4 females were group housed for two weeks, exposed to bedding from a male cage for 3 days, then individually exposed to a male for 24 h. Females were weighed on 6, 8, 10, and 12 days post coitum (dpc) to assess pregnancy status. The females that gained substantial body weight were considered pregnant (Heyne et al., 2015). Food, water and mice were weighed daily at 1000 h from 13 dpc to parturition. Treatments (CTL, HS and PF) were initiated on 14 dpc. On the day of delivery, we assessed litter size, litter mass, and pup survival rate. Pups were sacrificed by decapitation. Dams were sacrificed by decapitation under isoflurane anesthesia. Pair 2, 3, 4, and 5 mammary glands were dissected and weighed as previously described (Plante et al., 2011).

### Measuring lactogenesis of mammary glands

*Ex vivo* mammary gland lactogenesis was measured as previously reported with minor modifications (Mellenberger et al., 1973). Pair 4 and 5 mammary glands were sliced at an average thickness of 0.2 mm with a Thomas tissue slicer (Thomas Scientific, Swedesboro, NJ). Two slices of tissue from each gland were weighed and put into individual wells of a 24 well plate with 1 mL of Krebs-Ringer bicarbonate buffer without glucose. Slices were incubated at 37 °C with 5% CO_2_ for 0.5 h to allow for release of endogenous lactose within the tissue. Subsequently, tissue slices were transferred into another well containing 0.5 ml of Krebs-Ringer bicarbonate buffer supplemented with 10 mM glucose and 5 μg/ml of insulin (Sigma-Aldrich, St. Louis, MO) for a 3h incubation at 37 °C with 5% CO_2_. We collected media to assess lactose using a commercial assay kit (BioVision, Milpitas, CA). Lactose was corrected for tissue mass.

### Metabolic rates of mammary glands

We used a resazurin based assay that measures reducing equivalent production to assess mammary gland metabolic function (Beckett et al., 2018; Renquist et al., 2013). 1-3mg mammary tissue biopsies were isolated from mammary glands and weighed. Two biopsies from each gland were placed into 0.3 ml of pre-incubation medium (Dulbecco’s Modified Eagle’s Medium without glucose and phenol red (Bio5, University of Arizona, Tucson, AZ) supplemented with 1mg/ml BSA (Sigma-Aldrich), 0.1% DMSO and 1% Penicillin-Streptomycin solution (Thermo, Waltham, MA)) in a 96-well plate for 0.5h at 37 °C with 5% CO_2_. After a 30-minute pre-incubation, tissue biopsies were transferred into the assay medium for a 4h incubation (pre-incubation medium supplemented with 4 mg/ml glucose and 4.3% Alamar Blue solution (Bio-rad, Hercules, CA)). Fluorescence (excitation 530nm, emission 590nm) was measured at 0 and 4h of incubation in the assay medium. Change in fluorescence from 0 to 4h/mg tissue was calculated to understand tissue metabolic function.

### Statistics

We used the mixed procedure in SAS with main effect of treatment to analyze dependent variables that we had measured at a single timepoint, multiple comparisons were accounted for by Tukey’s adjustment (SAS Institute, Cary, NC). When appropriate, we performed repeated measures analyses with the dependent variables being treatment, day and their interaction. A Bonferroni correction was used to allow for multiple comparisons. Means were considered different when corrected P-value was less than 0.05. Means ± SEM were plotted using GraphPad PRISM (GraphPad Software, San Diego, CA).

## RESULTS

### Heat stress decreases food and water intake in adult male mice

We measured food and water intake of adult male mice while housed at the control environment for 4 days (CTL; 4d; 22 °C, relative humidity 50 %), during 5 days exposure to heat (HS; 5d; 35 °C, relative humidity 50 %), and again at CTL during a recovery period (CTL2; 3d). Heat exposure decreased food intake by 68.4% on the first day (P<0.0001; Fig. 1A) and heat continued to maintain food intake below that seen at CTL throughout the 5 days of heat exposure (*P*<0.05; Fig. 1A). Heat stress decreased average daily food intake by 46.5% (*P*<0.05; Fig. 1B). HS decreased dark cycle food intake throughout the 5 days (*P*<0.05), but only affected light cycle food intake on the first day of heat exposure (Figs 1C and 1D).

**Fig. 1.**
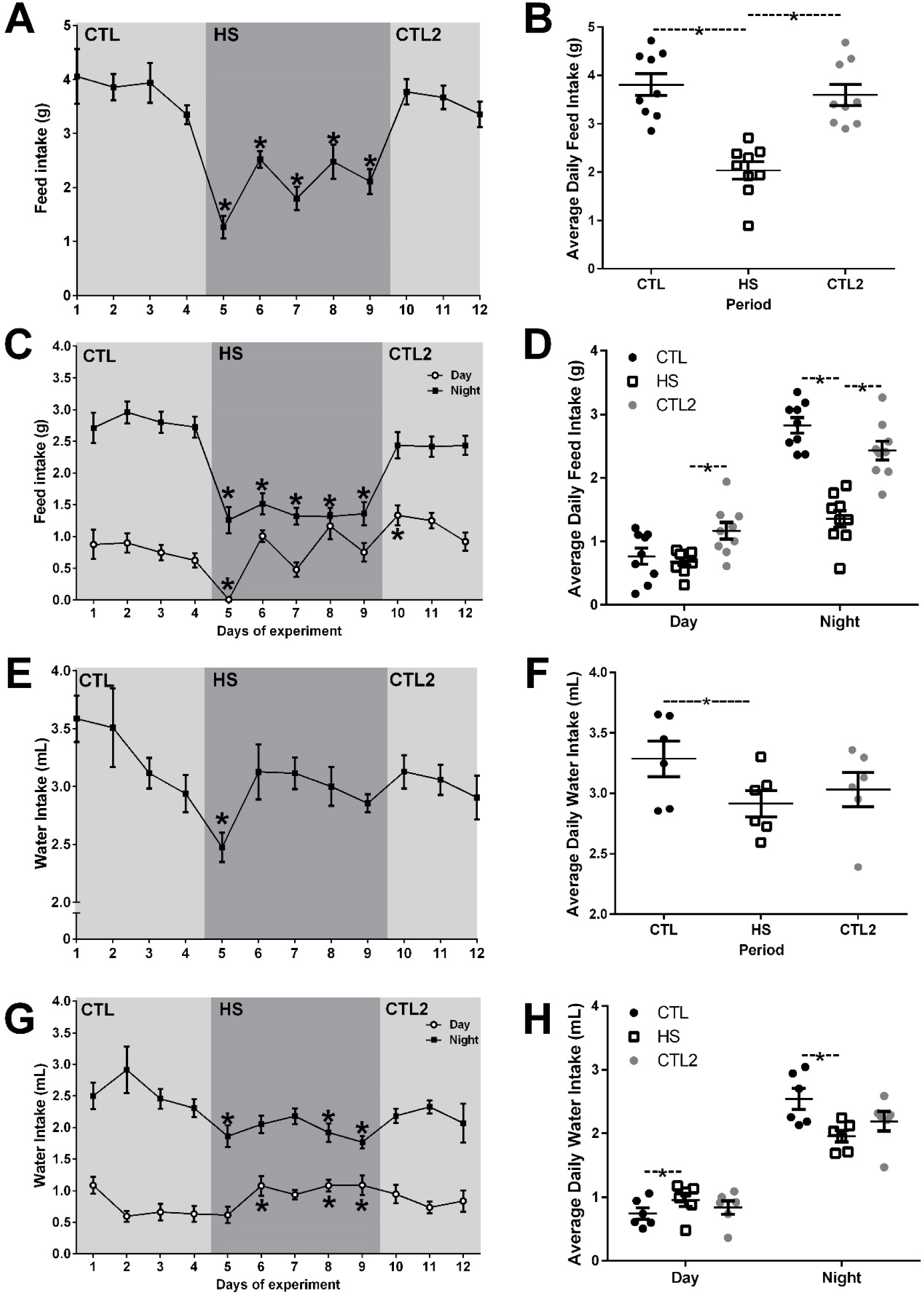
Food and water intake in 6 adult male mice (12-14 wks old) that were singly housed under control condition (CTL; 22°C, 50% humidity) from days 1-4 of the study, put in heat stress (HS; 35°C, 50% humidity) conditions from days 5-9, and returned to control condition (CTL2) from days 10-12. (A-B) 24h food intake, (C-D) light cycle and dark cycle food intake. (E-F) 24h water intake, (G-H) light cycle and dark cycle water intake. * indicates significant difference from mean of CTL (days 1-4; P < 0.05). -*-Indicates significant differences between indicated treatments (P < 0.05).

Heat exposure less robustly affected water intake, which decreased 24.7% on the first day of HS (*P*<0.05; Fig. 1E) and was not affected thereafter. Interestingly, heat did affect the diurnal pattern of water intake, decreasing water intake during the dark period and increasing water intake during the light period (*P*<0.05; Figs 1G and 1H).

### Heat exposure depresses body weight and food and water intake in lactating female mice

Heat exposure and pair-feeding decreased dam mass similarly, both decreased dam mass over the first 2 days of treatment that was maintained during the remainder of the study (*P*<0.05; Figs 2A and 2B). Although the decrease in body mass was maximal after 2 days of treatment, heat exposure suppressed food intake at all treatment days resulting in a cumulative food intake that was 41.6% lower than that of CTL dams (*P*<0.05; Figs 2C and 2D). Heat exposure similarly decreased water intake throughout the study, resulting in a cumulative 42.2% decrease in water consumption (*P* < 0.05; Figs 2E and 2F). Although pair-feeding resulted a similar depression of body weight and food intake, water intake was only mildly depressed by pair-feeding (11.1% cumulative decrease relative to control).

**Fig. 2.**
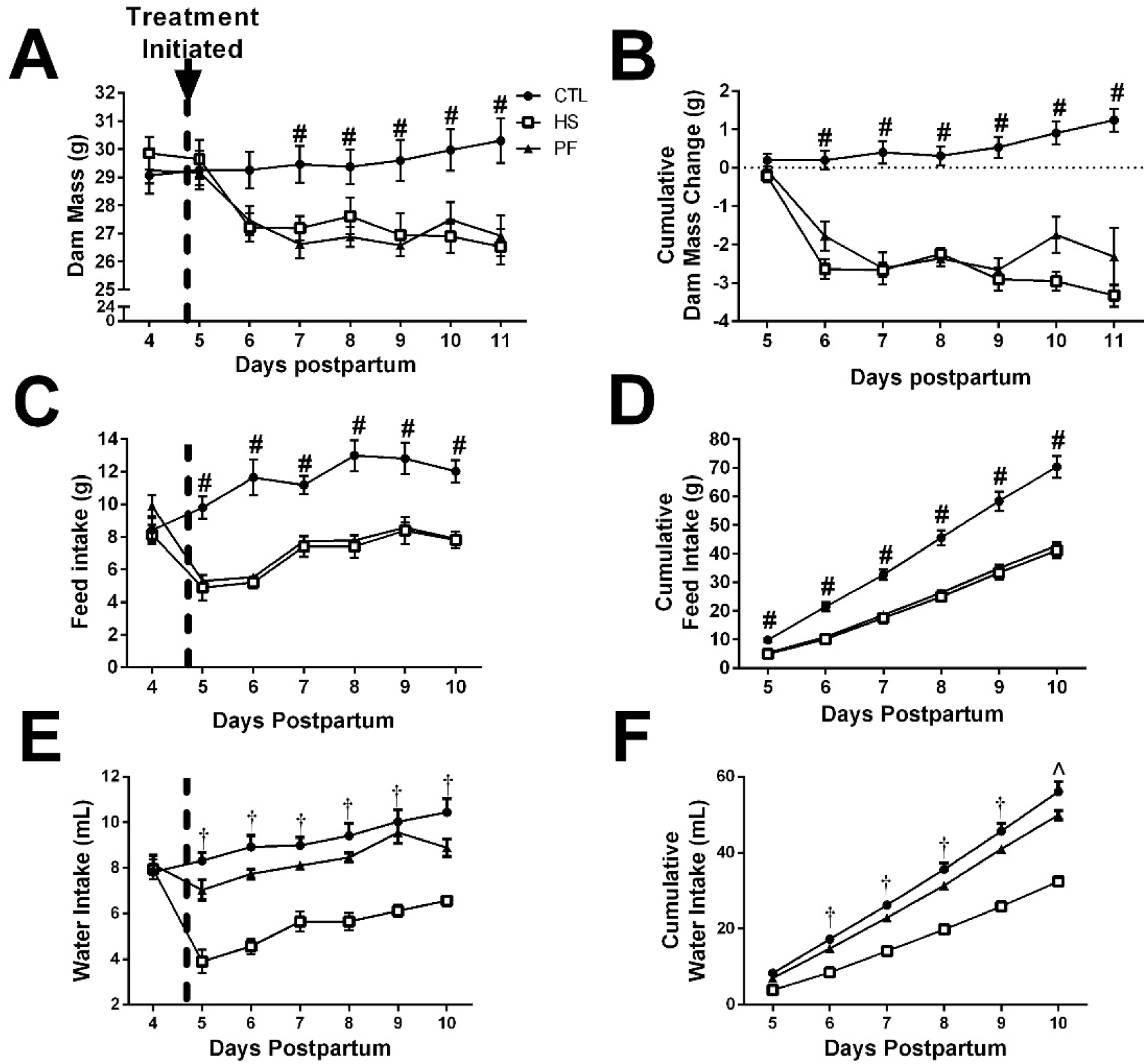
The response to heat stress (HS, n=9; 35°C, 50% humidity) and pair-feeding (PF, n=10; fed equivalent to HS mice) on from days 5-11 of lactation in multiparous dams on dam mass (A-B), food intake (C-D), and water intake (E-F). # control (CTL, n=10; 22°C, 50% humidity) > HS and PF groups, † indicates CTL and PF > HS, ^ indicates CTL > PF > HS (P < 0.05).

### Heat exposure during lactation depresses milk production through food intake dependent and independent mechanisms

We evaluated the effect of heat exposure on lactation performance by assessing litter mass and performing daily weigh suckle weigh measurements throughout the treatment duration (Fig. 3A-D). Within two days, heat exposure significantly decreased cumulative litter mass gain, which remained depressed throughout the treatment period (P < 0.05; Fig. 3B). After 6 days, litters from heat exposed dams had gained 35% less than those from control dams. Pair-feeding over those 6 days resulted in a 20% decrease in litter mass gain, with cumulative litter mass gain significantly differing from controls from days 4-6 of treatment. Accordingly, nearly 60% of the decrease in litter mass gain was explained by decreased food intake, while approximately 40% of the heat induced decrease in litter mass gain was independent of food intake. By using a weigh-suckle-weigh method, we were able to more directly assess milk production. A single day of heat exposure significantly decreased weigh-suckle-weigh litter mass change (P < 0.05; Fig. 3C). This heat-induced depression in litter mass weight gain during a suckling bout was maintained throughout the 6 days of heat exposure. As we previously observed with litter mass gain, pair-feeding resulted in daily weigh-suckle-weigh measures that were intermediate to those of control and heat-exposed mice. By expressing the change in mass that resulted from suckling as a cumulative measure across 6 days, we observed that heat stress decreased cumulative weigh-suckle-weigh mass change by 37.5% (Fig. 3D), very similar to the 35% decrease in total litter mass change (Fig. 3B). Pair-feeding resulted in a 17.5% decrease in cumulative weigh suckle weigh mass change (Fig. 3D), again very similar to the 20% decrease in total litter mass gain (Fig. 3B). Thus, 53.3% of the heat induced decrease in weigh suckle weigh mass change was independent the depression in food intake.

**Fig. 3.**
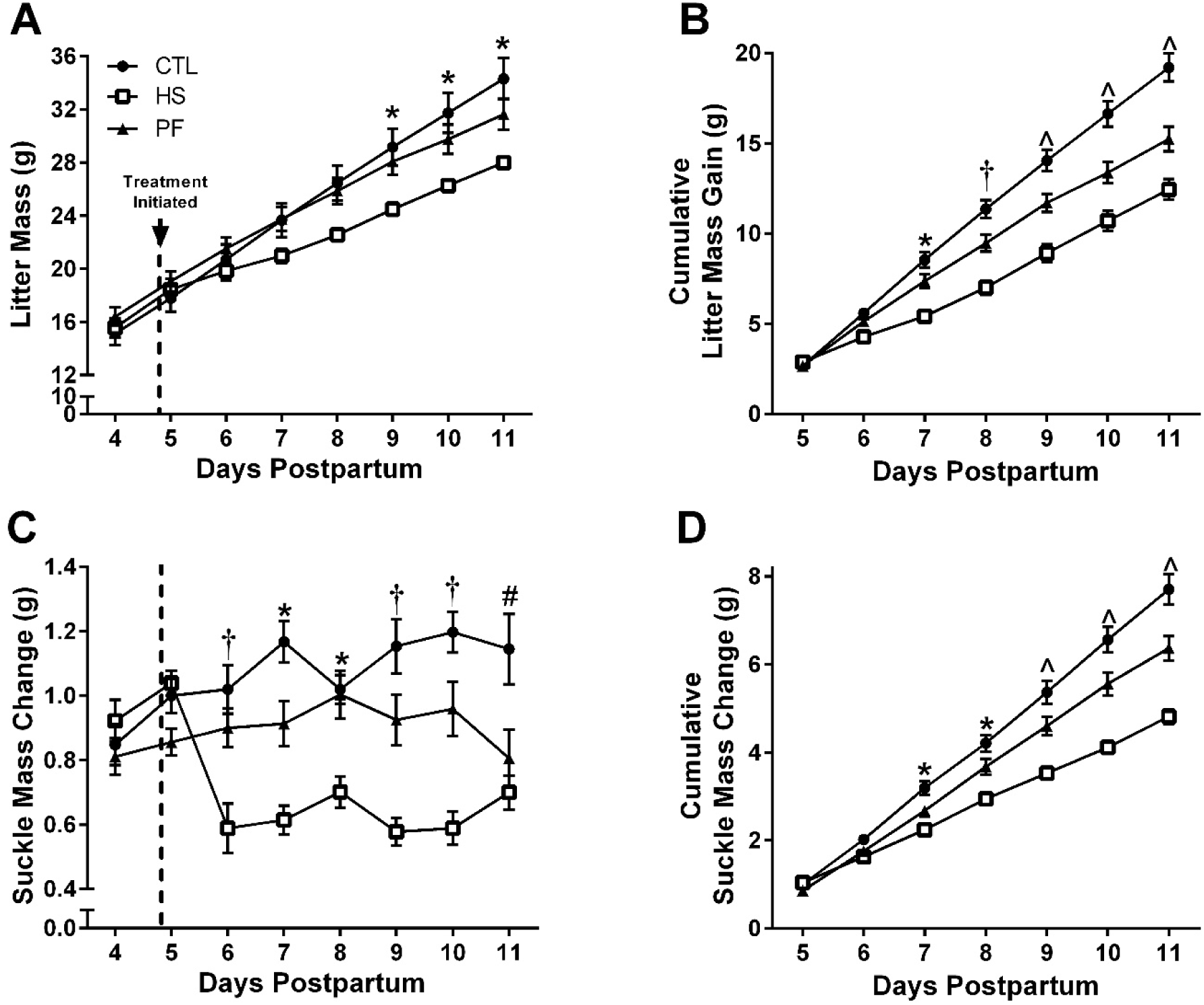
Litter mass (A-B) and lactation response (C-D) to heat stress (HS, n=9; 35°C, 50% humidity) or pair-feeding (PF, n=10; fed equivalent to HS mice) from days 5-11 postpartum. * indicates CTL (control, n=10; 22°C, 50% humidity) > HS, † indicates CTL and PF > HS, ^ indicates CTL > PF > HS, # CTL > PF and HS.

### Heat exposure during late gestation limits body weight gain and food intake without altering water consumption

Neither heat stress nor pair-feeding during late gestation affected dam mass (Fig. 4A). Surprisingly, pair-feeding more robustly decreased dam mass gain during late gestation (34.9%) than did heat stress (18.1%; Fig. 4B).

**Fig. 4.**
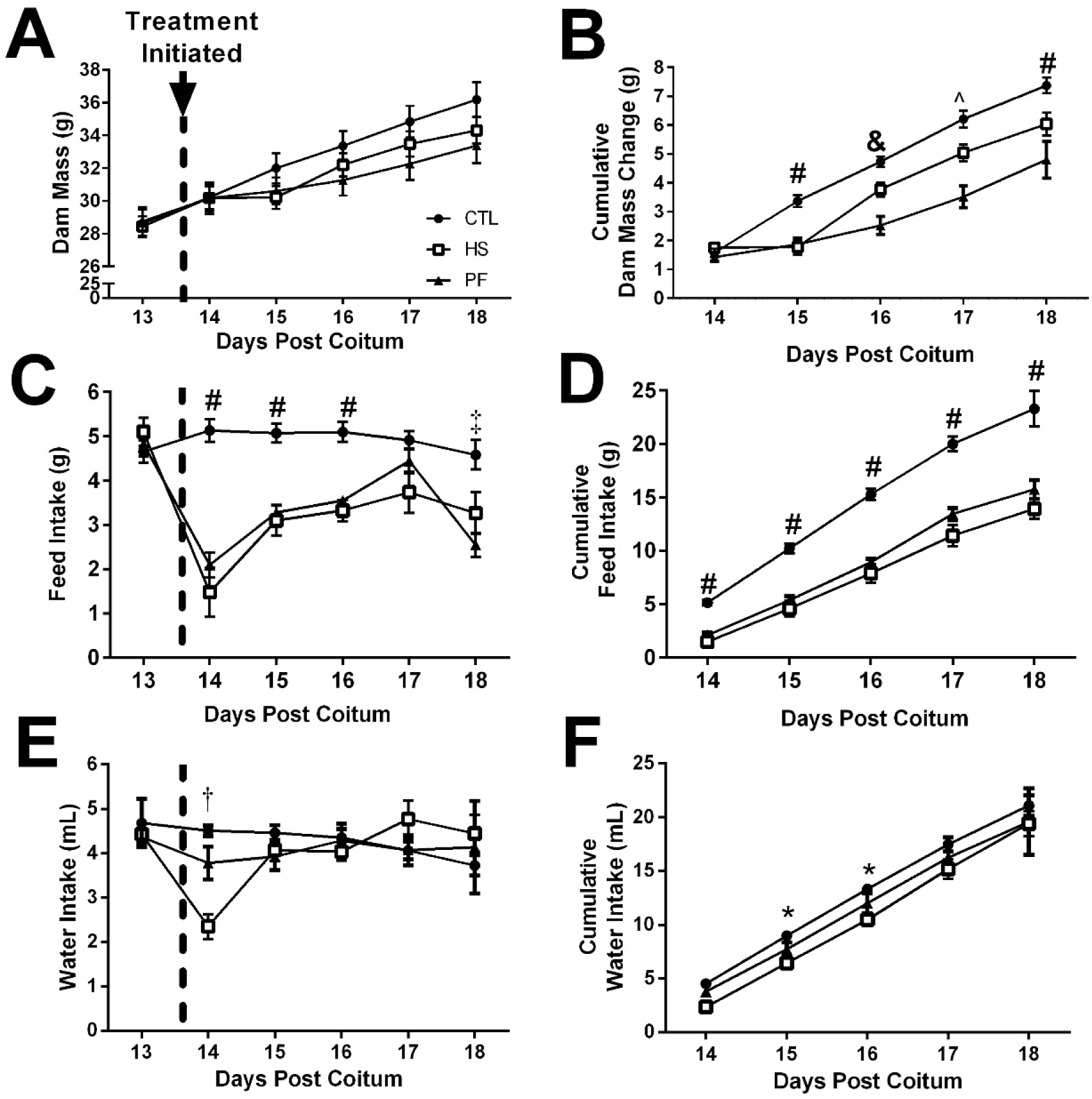
The effect of heat stress (HS, n=10; 35°C, 50% humidity) and pair-feeding (PF, n=10; fed equivalent to HS mice) from 14 days post coitum to parturition on dam mass (A-B), food intake (C-D), and water intake (E-F). * indicates control (CTL, n=11; 22°C, 50% humidity) > HS, † indicates CTL and PF > HS, # indicates CTL > PF and HS, ^ indicates CTL and HS > PF, & indicates CTL and HS > PF, ‡ indicates CTL > PF.

Heat exposure decreased daily food intake (Fig. 4C) on 4 out of 5 days and cumulative food intake throughout the entire study (P < 0.05; Fig. 4D). In fact, heat exposure decreased cumulative food intake by nearly 40%. Surprisingly, heat exposure only decreased water intake on the first day of treatment (P < 0.05; Fig. 4E). Since heat did not affect water intake after the first day of exposure there was no effect of heat on cumulative water intake in the end (Fig. 4F). Pair-feeding late gestation dams did not alter daily or cumulative water intake (Figs. 4E and 4F).

### Heat exposure during late gestation affects litter viability and mean pup mass without altering gestation length, litter size, or litter mass

Gestation length, litter size and litter mass at birth were not affected by either heat stress or pair-feeding (Figs 5A, 5B and 5D). However, dam heat exposure decreased pup survival (60%) and mean pup mass (16 %; *P*<0.05; Figs. 5C and 5E). These effects on survival and pup mass are independent of food intake, as pair-feeding did not affect either variable.

**Fig. 5.**
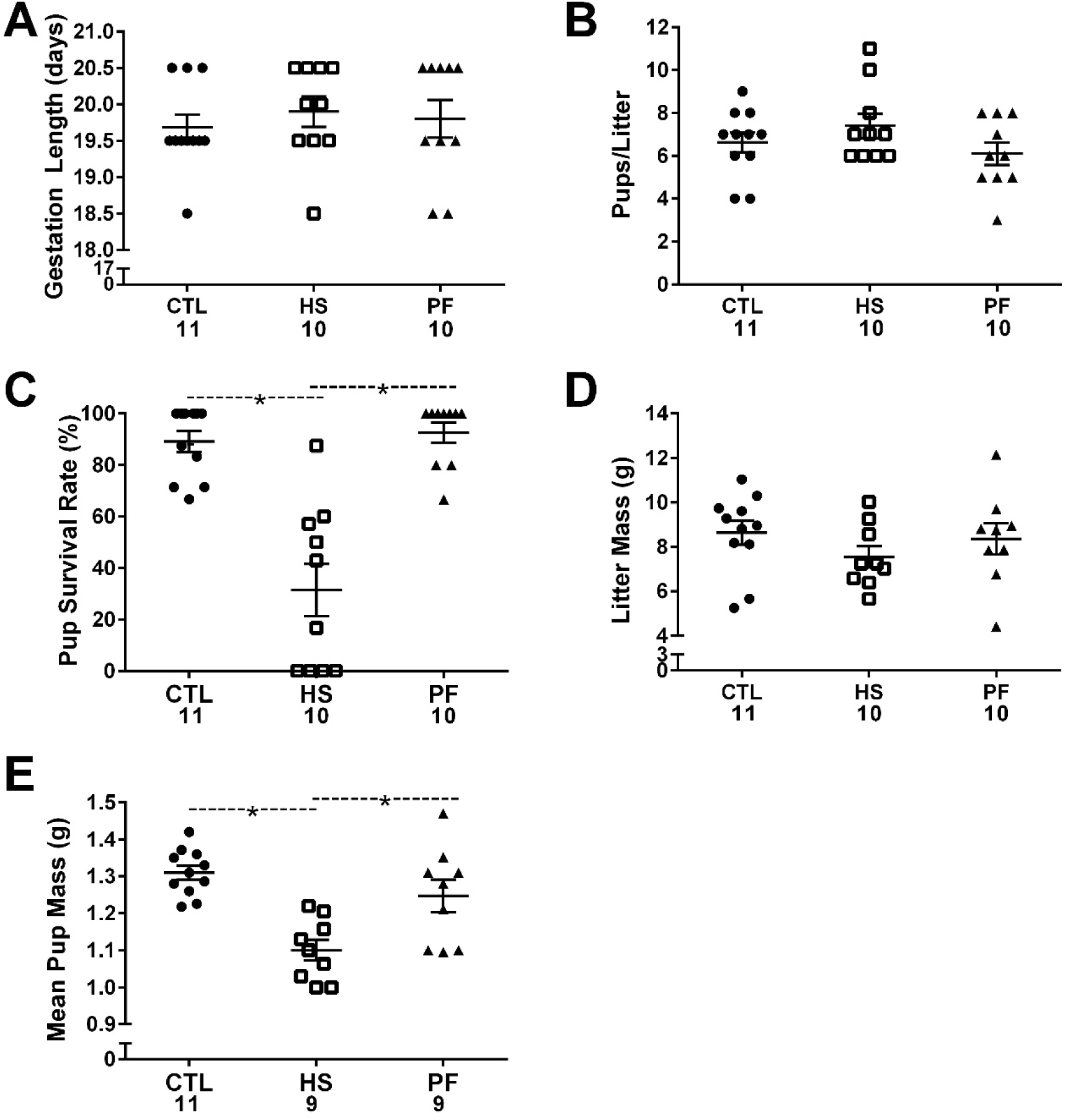
The effect of heat stress (HS; 35°C, 50% humidity) and pair-feeding (PF; fed equivalent to HS mice) of virgin female mice from 14 days post coitum to parturition on A) gestation length, B) pups/litter, C) pup survival rate, D) litter mass, and E) mean pup mass. Numbers under the treatment donate number of biological replicates. *indicates significant differences between treatments (P < 0.05).

### Heat exposure during late gestation depresses mammary gland mass and function

Heat stress and pair-feeding similarly depressed mammary gland mass at parturition (P<0.05; Fig. 6A). Heat exposure during late gestation depressed e*x vivo* reducing equivalent production per mg mammary tissue, while there was no effect of pair-feeding (P < 0.05; Fig. 6B). Heat exposure during the last 5-6 days of gestation decreased *ex vivo* lactose production independent of the decrease in food intake (P < 0.05; Fig. 6C and 6D). In fact, pair-feeding did not affect either the lactose production/mg tissue or lactose production/gland. This data proposes that heat exposure decreases mammary gland mass dependent on decreased food intake, while affecting mammary function through a food intake independent mechanism.

**Fig. 6.**
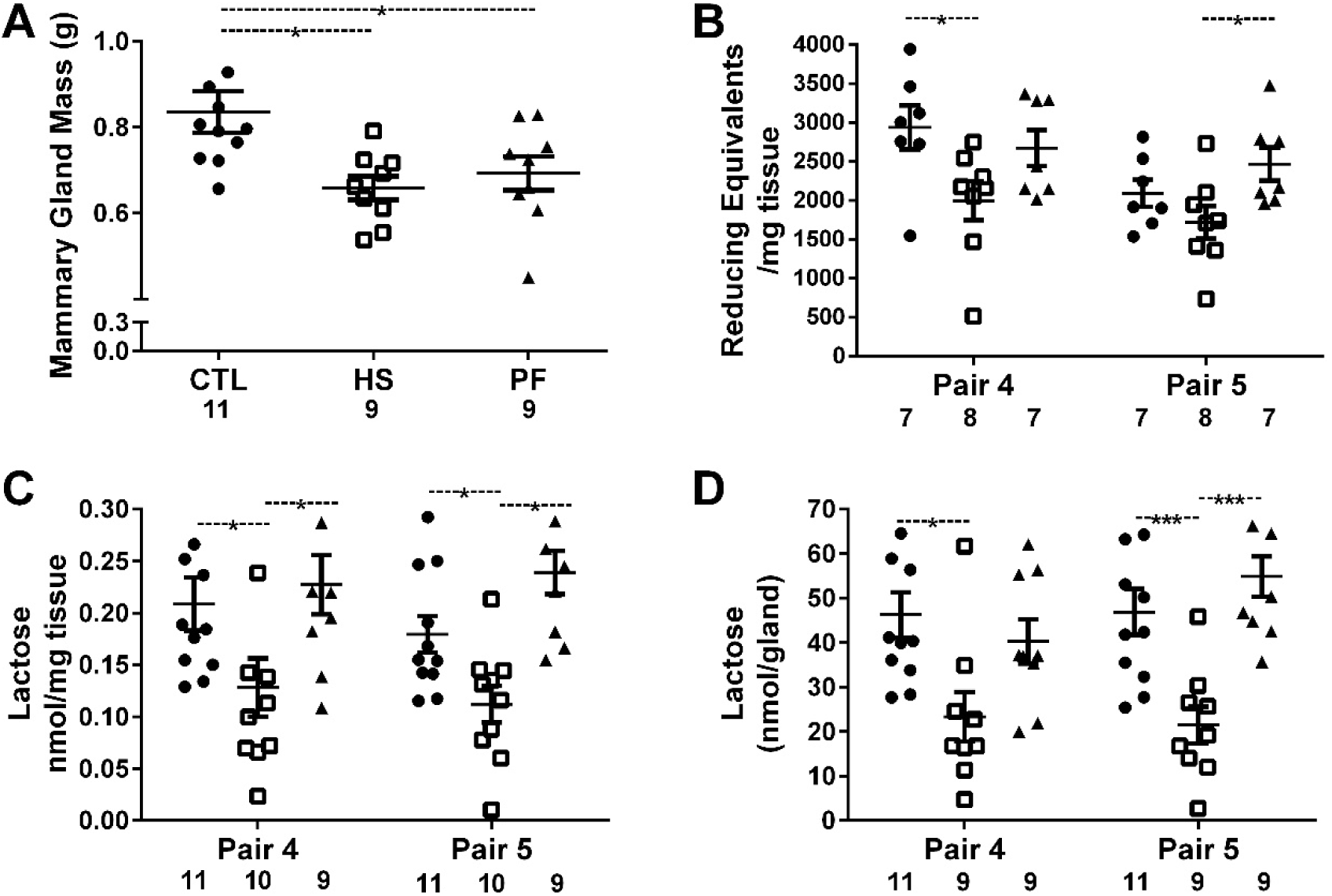
The effect of heat stress (HS; 35°C, 50% humidity) and pair-feeding (PF; fed equivalent to HS mice) from 14 days post coitum to parturition on A) mammary gland mass, B) metabolic rate, C) lactogenesis ex vivo, D) predicted lactose production per gland. Numbers under the treatment donate number of biological replicates. *indicates significant differences between treatments (P < 0.05), *** P < 0.001.

## DISCUSSION

The hypophagic response to heat exposure is conserved across homeothermic animals, decreasing growth and product synthesis (milk, eggs) (Barrett et al., 2019; O’Brien et al., 2010; Rhoads et al., 2009; Zhao et al., 2018). We aimed to understand the hypophagia dependent and independent effects of heat exposure during peak lactation and late gestation on milk production and mammary gland development, respectively.

### Relationship between hypophagia and hypogalactia under heat stress conditions

In lactating mice, the energetic demands of lactation are robust. In fact, 44% of gross energy intake is used for milk production (Johnson et al., 2001). To meet the energetic demand for lactation, lactating dam food intake is nearly 3 times higher than in male mice of similar mass and 2 time higher than in the late gestation dam. Similarly robust effects of lactation on food intake have been reported in the C57Bl6/J mouse strain used here (Makarova et al., 2010). With this increased metabolic demand our lactating dams recapitulate the increased metabolic demand in the lactating cow and the resulting increased sensitivity to exogenous heat (Collier et al., 2012). As evidence, lactation rises body temperature by 1.1°C in mice maintained at 21°C causing chronic hyperthermia (Gamo et al., 2013).

Reduced food intake decreases the milk production across species. Consuming 32% less energy (less than 1500 kcal/d) for a week caused 15% depression in milk production in women (Strode et al., 1986). In rats, 50% food intake restriction leads to a 66% decrease in milk yield (Warman and Rasmussen, 1983). Given that heat depresses food intake, we aimed to understand the role of heat induced hypophagia in the depression of milk production. By using litter mass gain and weigh suckle weigh measures as proxies for milk production, we showed that approximately 50% of heat induced hypogalactia was independent on the heat induced depression in food intake. Our findings recapitulate findings in the dairy cow which have established a nearly identical relative role for hypophagia (50%) in heat induced hypogalactia (Rhoads et al., 2009; Wheelock et al., 2010).

### Heat exposure induced food intake depression impairs mammary gland weight but not mammary function and fetal growth

From late gestation through early lactation there is robust mammary gland expansion through proliferation (Howard and Gusterson, 2000; Knight and Peaker, 1982; Lu and Anderson, 1973; Sorensen et al., 2002). Late gestation heat stress limits mammary gland development, depressing milk production throughout lactation (Collier et al., 1982; Hooper et al., 2019; Tao et al., 2011). This depression in milk production that lasts the entire lactation is economically disastrous.

Heat exposure during late gestation decreased dam weight gain (Fig. 4A) as previously observed in the dairy cow (Collier et al., 1982; Tao et al., 2011). We found gestational heat exposure caused a 20% reduction in mammary gland weight at parturition. Indeed, mammary cells proliferation, but not apoptosis, is vulnerable to thermal stress during pregnancy (Tao et al., 2011). Food intake and body weight gain during pregnancy are both associated with mammary gland DNA content (Kumaresan and Turner, 1968). We found that pair-feeding recapitulated the impaired mammary growth that we observed in heat-exposed animals (Fig. 6A). Mammary gland function measured as mammary gland mitochondrial activity and lactose production were impaired by heat exposure (Fig. 6B-6C). Pair-feeding did not recapitulate the heat-induced loss in function. Similarly, in the cow, heat stress decreases mRNA expression of genes involved in production of key milk proteins (casein and lactalbumin) and transport of amino acids and glucose (Gao et al., 2019) when compared to that observed in pair-fed cows. In cows, heat exposure also appears to limit the ability of the mammary gland to mobilize fatty acids from triglycerides and catabolize those fatty acid through β-oxidation (Adin et al., 2009; Gao et al., 2019). Accordingly, depressed mammary function appears to be a direct response to heat and may be a result of limited metabolite flux.

Late gestation heat stress has been reported to either shorten or not affect gestation length in human and farm animals (Collier et al., 1982; Porter et al., 1999; Tao et al., 2012; Williams et al., 2013). We found no effect of gestational heat stress on gestation length in mice. However, we did observe that late gestation heat stress depressed neo-natal mass and survival. In fact, the degree of decrease we observed (16%) is in the range (6% ∼ 30%) of that observed in ruminant species (Laporta et al., 2017; Tao and Dahl, 2013). Gestational heat stress caused malfunction of placenta and decreases placental blood flow, in turn resulting in depressed fetal growth (Alexander et al., 1987; Reynolds et al., 1985). In mice, the stress hormone corticosterone reduced blood vessel density in the placenta, which led to fetal growth restriction (Vaughan et al., 2012). Since we observed normal fetal growth in pair-fed dams, nutrient accessibility to fetus rather than nutrient intake by the dam likely contributed to the fetal restriction. Depressed fetal development may negatively impact subsequent performance of the offspring. In fact, *in utero* heat stress has been shown to depress milk production and mammary gland structure in offspring’s first lactation (Fabris et al., 2019; Monteiro et al., 2016; Skibiel et al., 2018).

Our studies establish that the mouse recapitulates the heat stress phenotypes observed in production animals. These include depressed food intake, decreased lactation that is both dependent and independent of feed intake, and depressed mammary gland development. Using this mouse model, we observed the novel finding that heat-induced depression of mammary gland mass was completely attributable to depressed feed intake, while the decreased mammary function was entirely independent of hypophagia. Our late gestation heat exposure caused restricted fetal growth and reduced liveborn rate, which is similar to that observed in farm animals (Laporta et al., 2017; Monteiro et al., 2016; Tao and Dahl, 2013). These in utero effects were independent of changes in food consumption. Together, our data validate these mouse models as valuable tools for studying the physiological responses to heat stress. Thereby opening the door for mechanistic studies using genetic and pharmacologic models to identify the mechanism by which heat exposure causes these physiological changes.

